# A Nasal Spray Solution of Grapefruit Seed Extract plus Xylitol Displays Virucidal Activity Against SARS-Cov-2 *In Vitro*

**DOI:** 10.1101/2020.11.23.394114

**Authors:** Gustavo Ferrer, Arian Betancourt, Camille Celeste Go, Hector Vazquez, Jonna B. Westover, Valeria Cagno, Caroline Tapparel, Marcos A. Sanchez-Gonzalez

## Abstract

The severe acute respiratory syndrome coronavirus 2 (SARS-CoV-2), responsible for the ongoing pandemic coronavirus disease 2019 (COVID-19) has triggered worldwide concerted efforts in an attempt to identify effective therapies. In the present study, we have identified two candidate agents with potential activity against SARS-CoV-2 which can be administered intranasally, namely, xylitol and grape seed fruit extract (GSE). A commercially available nasal spray (Xlear) combining xylitol and GSE has been available for years, but the antiviral effects of this solution have not been documented. This *in vitro* study examined the virucidal effect of Xlear against SARS-CoV-2. To this end, two independent sets of experiments were carried out to test the hypothesis that Xlear is an effective (Experiment I) and replicable (Experiment II) means to deactivate SARS-CoV-2. When tested against SARS-CoV-2, the test compound GSE 0.2% was the only compound effective at reducing >3 log10 CCID50 infectious virus from, 3.67 log10 CCID50/0.1 mL to an undetectable amount of infectious virus. The present results validated by two independent sets of experiments, performed by different labs, on different viral strains, provide early evidence to encourage further pilot and clinical studies aimed at investigating the use of Xlear as a potential treatment for COVID-19

## 1 Introduction

The initial global outbreak of the severe acute respiratory syndrome coronavirus 2 (SARS-CoV-2), responsible for the ongoing pandemic coronavirus disease 2019 (COVID-19), was initially identified in Wuhan, China in December 2019. As of July 2020, there were more than 13.3 million confirmed cases worldwide, with total deaths exceeding 573,000 (Dong et al., 2020). Worldwide concerted efforts have been made in an attempt to characterize the disease and identify effective therapies targeting SARS-CoV-2 including lines of studies focusing on the route of infection, the potential routes of administration of therapeutic agents as well as the potential efficacy of antiseptics (Meister et al., 2020). In this vein, a landmark study found that the coronavirus infects the nasal cavity via the angiotensin-converting enzyme 2 (ACE2) protein which appears to be the host-cell receptor for SARS-CoV-2 (Hoffmann et al., 2020). Since the nasal epithelium cells have the highest percentage of ACE2 expressing ciliate cells in the proximal airways, it is plausible to suggest that pharmacological agents such as sprays that are used via the intranasal route of administration might be optimal candidates for providing effective therapies against COVID-19 (Jia et al., 2005).

In a recent literature review conducted by Higgins et al. it is highlighted that intranasal drug delivery represents an important area of research for viral diseases and COVID-19 (Higgins et al., 2020). They concluded that the intranasal method of drug delivery has potential relevance for future clinical trials in the setting of disease prevention and treatment of SARS-CoV-2 in addition to other viral diseases (Higgins et al., 2020). Subsequently, Siddiqi et.al (2020), in a diagram of COVID-19 disease progression, illustrated that the viral response phase is highest during the early infection of the disease process, of which patients manifest mild constitutional symptoms. Taken together the aforementioned studies support our rationale that therapeutic strategies should be aimed at reducing the viral load in the nose by targeting this mild-moderate phase of the disease process, and hence the use of a nasal spray might be an effective means to accomplish this therapeutic strategy.

In the present study, we have identified two candidate agents with potential activity against SARS-CoV-2 which can be administered intranasally, namely, xylitol and grape seed fruit extract (GSE). Xylitol, a sweetener with antimicrobial and anti-inflammatory properties, has been shown effective in decreasing the incidence of dental caries and improving chronic rhinitis as well as important microbiota and immunological modulatory effects (Akgül et al., 2020; Haukioja et al., 2008; Weissman et al., 2011; Xu et al., 2016). Xylitol has been reported to have multiple health benefits as well as is generally safe and well-tolerated for most adults in doses up to 35 grams per day and up to 20 grams per day in children (Salli et al., 2019; Storey et al., 2007; Ur-Rehman et al., 2015). A derivative of grapefruit seeds, GSE, is associated with abundant health benefits due to the presence of antioxidants and proanthocyanidin complexes (Chacón et al., 2009). Also, GSE has been documented to have inhibitory effects against the avian influenza virus, Newcastle disease virus, infections bursal disease virus, as well as other pathogenic enteric viruses (Komura et al., 2019; Su and D’Souza, 2011). A commercially available nasal spray combining xylitol and GSE, marketed as Xlear (American Fork, UT, USA), has been widely used in the United States for several decades, but the antiviral effects of this solution have not been documented. Accordingly, the aim of the present *in vitro* study was to examine the virucidal effect of Xlear against SARS-CoV-2. To this end, two independent sets of experiments were carried out to test the hypothesis that Xlear is an effective (Experiment I) and replicable (Experiment II) means to deactivate SARS-CoV-2 the causative microorganism of COVID-19.

## 2 MATERIALS AND METHODS

### 2.1 Experiment I: Xlear Virucidal Activity Efficacy

#### 2.1.1 Procedure

SARS-CoV-2, USA-WA1/2020 strain, virus stock was prepared before testing by growing 2 passages in Vero 76 cells. Culture media for prepared stock (test media) was MEM with 2% fetal bovine serum (FBS) and 50 µg/mL gentamicin. Human rhinovirus 16, strain 11757 purchased from ATCC (Gaithersburg, Maryland, USA), was grown in 3 passages of HeLa cells in MEM with 2% fetal bovine serum (FBS), 25 mM MgCl2, and 50 µg/mL gentamicin. Test media is the growth media with 5% FBS.

#### 2.1.2 Virucidal Assay

Test compounds including commercially available Xlear containing purified water, 11% Pure Xylitol (Shandon Lujian, Shandong, China), 0.6%NaCL (Saline), and 0.015% GSE (Chemie Research & Manufacturing Co., Casselberry, FL, USA) were obtained from the manufacturer in liquid form and stored at room temperature. The test compound 11% xylitol in saline was diluted 1:2 with water before testing. Each solution was mixed directly with virus stock so that the final concentration was 90% of each test compound and 10% virus stock. A single concentration was tested in triplicate. Test media without virus was added to duplicate tubes of the compounds to serve as toxicity and neutralization controls. Ethanol (90%) was tested in parallel as a positive control and water only as a virus control. The test solutions were incubated at room temperature (22 ± 2°C) for 15 minutes with SARS-CoV-2 or Rhinovirus-16. The solutions were then neutralized by a 1/10 dilution in the test media of each specific virus. The virucidal assays were performed in triplicate, then after neutralization, the triplicate samples were pooled, serially diluted, and assayed for infectious virus.

#### 2.1.3 Virus Quantification

The surviving virus from each sample was quantified by standard end-point dilution assay. Briefly, the neutralized samples were pooled and serially diluted using eight log dilutions in test medium. Then 100 µL of each dilution was plated into quadruplicate wells of 96-well plates containing 80-90% confluent Vero 76 (SARS-CoV-2) or HeLa cells (Rhino-16). The toxicity controls were added to an additional 4 wells of Vero 76 or HeLa cells and 2 of those wells at each dilution were infected with virus to serve as neutralization controls, ensuring that the residual sample in the titer assay plate did not inhibit growth and detection of the surviving virus. Plates were incubated at 37 ± 2°C with 5% CO_2_ for 5 days and at 33 ± 2°C with 5% CO2 for 4 days for the SARS-CoV-2 assay and the Rhinovirus-16 assay, respectively. Each well was then scored for the presence or absence of an infectious virus. The titers were measured using a standard endpoint dilution 50% cell culture infectious dose (CCID50) assay calculated using the Reed-Muench (1948) equation and the log reduction value (LRV) of each compound compared to the negative (water) control was calculated.

### 2.2 Experiment II: Xlear Virucidal Activity Replication

#### 2.2.1 Procedure

SARS-CoV2/Switzerland/GE9586/2020 virus stock was amplified and titrated in Vero E6 cells by plaque assay cultured in DMEM HG with 5% fetal bovine serum (FBS) and 1% penicillin/streptomycin.

##### Dose-response Assay

Xlear nasal spray was serially diluted in DMEM HG and incubated with SARS-CoV2 (MOI 0.003 corresponding to 200 pfu/well) for 1 hour at 37°C and subsequently added on Vero E6 cells for 1 hour at 37°C. The inoculum was then removed, cells were washed and overlaid with DMEM HG with 5% FBS and Avicel 0.8%. 48hpi cells were fixed with PFA 4% and stained with crystal violet. Plaques were counted and percent of infection calculated in comparison with untreated wells. The experiments were performed twice independently, and each was performed in duplicate.

##### Virucidal Assay

Xlear spray was mixed in different concentrations with SARS-CoV2 stock (10^5^pfu). The compound was mixed directly with the virus solution with a final concentration of respectively 90%, 80%, 60%, or 20%. PBS was used as control. The solution and virus were incubated at 37 °C for 1 hour. The solution was then neutralized by a 1/10 dilution in test media. A 60% condition was repeated in two independent experiments while the other dilutions were performed in a single experiment in duplicate.

The infectious virus from each sample was quantified by standard end-point dilution assay. 100 µL of each dilution were plated into quadruplicate wells of 96-well plates containing 80-90% confluent Vero 76 cells. Plates were incubated at 37°C with 5% CO2 for three days. Each well was then scored for the presence or absence of the virus. The end-point titers (TCID50) values were calculated using the Reed-Muench (1948) equation.

#### 2.2.2 Toxicity assay

Vero-E6 (13000 cells per well) were seeded in 96-well plate. Xlear was serially diluted in DMEM supplemented with 5% FBS and added on cells for 1h, followed by a washout, addition of DMEM supplemented with 5% FBS for additional 48h hours. MTT reagent (Sigma Aldrich) was added on cells for 3h at 37°C according to manufacturer instructions, subsequently cells were lysed with pure DMSO and absorbance read at 570 nm. Percentages of viability were calculated by comparing the absorbance in treated wells and untreated.

## 3 RESULTS

### 3.1 Experiment I

Virus titers and LRV of Rhinovirus-16 and SARS-CoV-2 when incubated with a single concentration of the Xlear solutions are shown in Table 1. After a 15-minute contact time, the Xlear nasal spray was not effective at reducing the infectious Rhino-16 virus. When tested against SARS-CoV-2, the test compound GSE 0.2% was the only compound effective at reducing >3 log10 CCID50 infectious virus from, 3.67 log10 CCID50/0.1 mL to an undetectable amount of infectious virus (Table 1). The Xlear nasal spray and the GSE 0.2% had some toxicity in the top rows (1/10 dilution of the test sample) which may have contributed to the virucidal effect of the GSE. The 11% xylitol and 11% erythritol had no cytotoxicity. The positive control and neutralization control performed as expected.

**Table 1.**
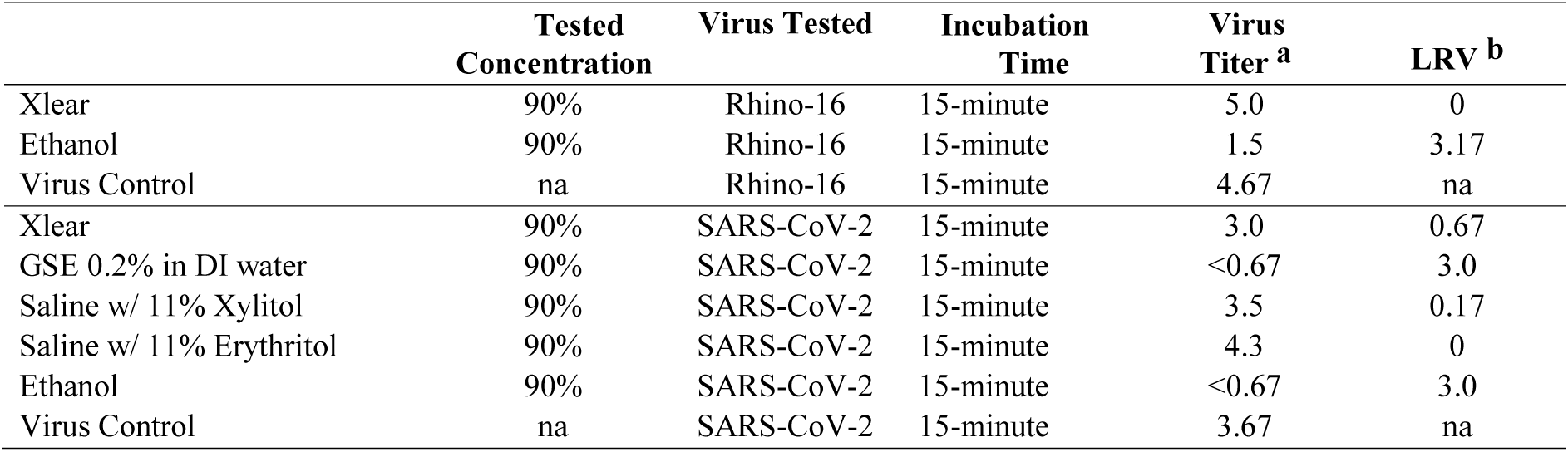
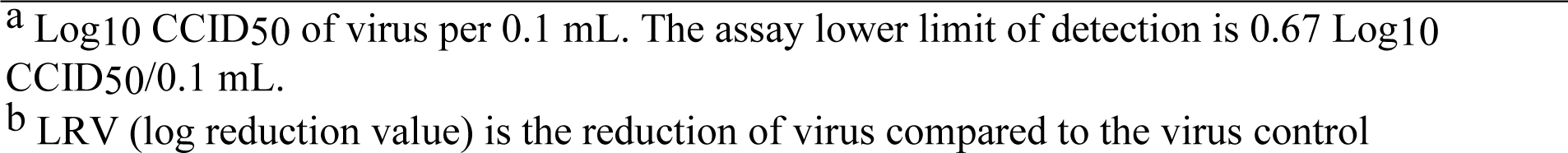
Virucidal efficacy of Xlear compounds against Rhinovirus-16 and SARS-CoV-2 after a 15-minute incubation with virus at 22 ± 2°C.

|  | **Tested**<br>**Concentration** | **Virus Tested** | **Incubation**<br>**Time** | **Virus**<br>**Titer <sup>a</sup>** | **LRV <sup>b</sup>** |
| --- | --- | --- | --- | --- | --- |
| Xlear | 90% | Rhino-16 | 15-minute | 5.0 | 0 |
| Ethanol | 90% | Rhino-16 | 15-minute | 1.5 | 3.17 |
| Virus Control | na | Rhino-16 | 15-minute | 4.67 | na |
| Xlear | 90% | SARS-CoV-2 | 15-minute | 3.0 | 0.67 |
| GSE 0.2% in DI water | 90% | SARS-CoV-2 | 15-minute | <0.67 | 3.0 |
| Saline w/ 11% Xylitol | 90% | SARS-CoV-2 | 15-minute | 3.5 | 0.17 |
| Saline w/ 11% Erythritol | 90% | SARS-CoV-2 | 15-minute | 4.3 | 0 |
| Ethanol | 90% | SARS-CoV-2 | 15-minute | <0.67 | 3.0 |
| Virus Control | na | SARS-CoV-2 | 15-minute | 3.67 | na |

### 3.2 Experiment II

SARS-Cov2 is inhibited in the dose-response assay (Figure 1) by different concentrations of Xlear spray. However, the dilution 1:2 in medium evidenced damage to the cells with almost complete loss of the cells, while with the dilution 1:6 a partial damage to the cell was evidenced, while no morphologic changes in cells were visible from dilution 1:12 onwards. These results were further confirmed with toxicity assays (Figure 1b).

**Figure 1.**
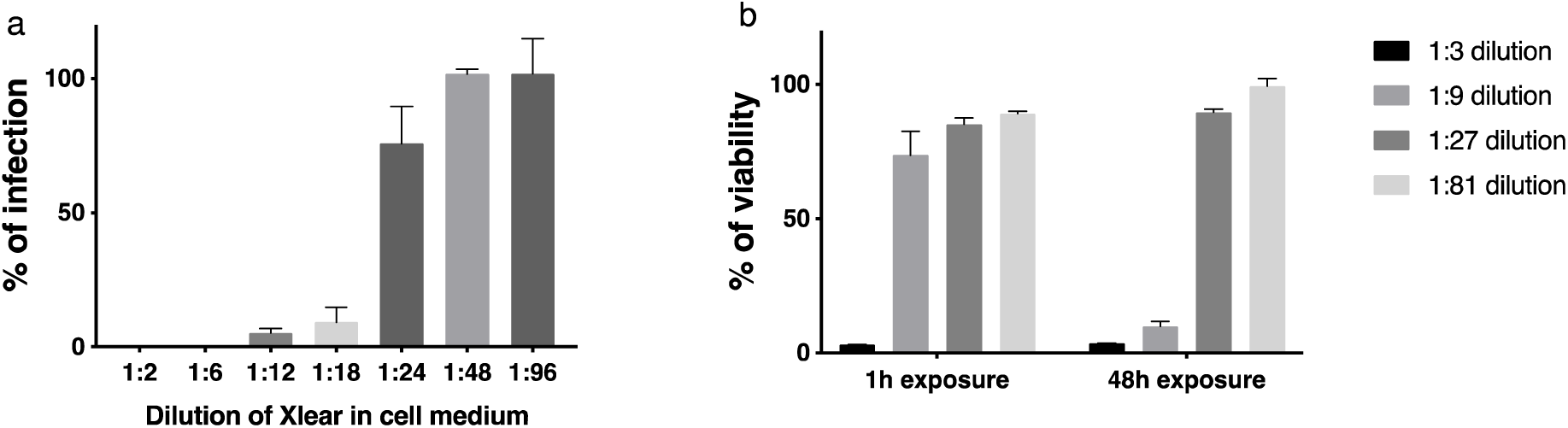
a) SARS-CoV-2 dose-response inhibition. Xlear was incubated at different dilutions with SARS-CoV2 (200 pfu) for 1h at 37 C. At the end of the incubation, mixtures were serially diluted and added for 1h at 37°C on Vero-E6 cells. Mixtures were then removed, and cells overlaid with medium containing 0.8% avicel. Cells were fixed 48hpi and plaques were counted. Results are mean and SEM of 2 independent experiments performed in duplicate. b) Xlear toxicity evaluation. Different dilutions of the nasal spray were incubated for 1h (followed by addition of medium for 47h) or for 48h on cells in DMEM 5% FBS. At the end of the incubation MTT reagent was added on cells and percentages of viability were evaluated by comparing treated and untreated wells.

In the virucidal assays (Figure 2), Xlear showed virucidal activity at the different concentrations tested. Complete inhibition of viral infectivity was observed for the 90%, 80%, 60% condition, and a reduction of 2.17 log of viral titer in the 20% condition. In this assay, the mixture of virus and Xlear was neutralized by a 1/10 dilution before addition on cells, therefore diluting the compound below the toxic doses determined in the toxicity assay (Figure 1b).

**Figure 2.**
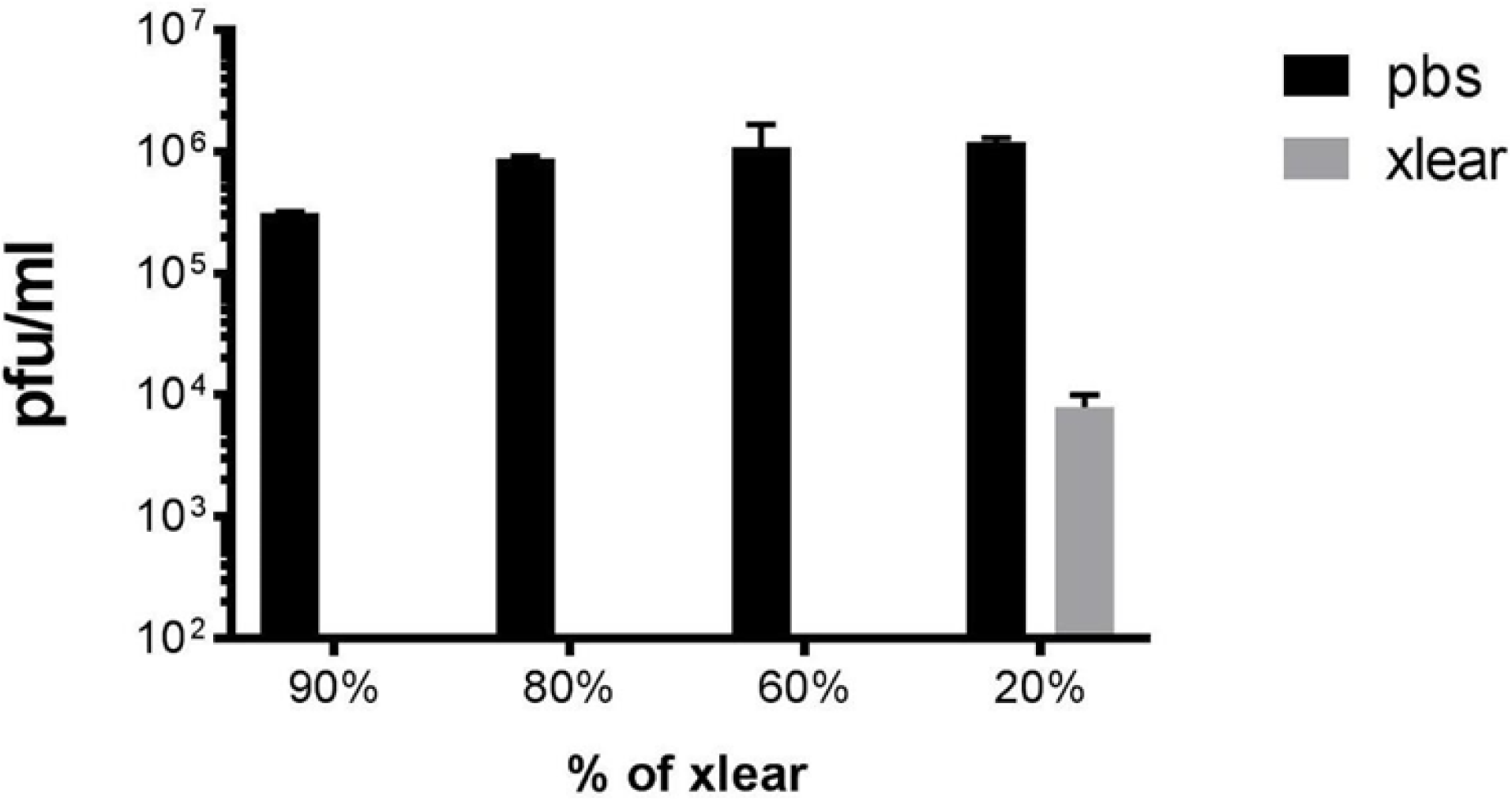
SARS CoV-2 virucidal assay. Xlear was incubated with SARS-CoV2 (5*10^5^ pfu) for 1h at 37 C. At the end of the incubation, mixtures were serially diluted and added on Vero-E6 cells. Cells were fixed 48hpi and scored for presence or absence of cytopathic effect and TCID50/ml was determined. Results are mean and SD of two independent experiments.

## 4 DISCUSSION

The present study sought to evaluate the *in vitro* virucidal effects of a solution combining xylitol and GSE in a nasal spray formulation known as Xlear. The novel results of this study support our hypothesis that Xlear displays virucidal activity against SARS-CoV-2. The present results validated by two independent sets of experiments, performed by different labs, on different viral strains, provide early evidence to encourage further pilot and clinical studies aimed at investigating the use of Xlear as a potential treatment for COVID-19.

Xlear is a solution of xylitol and GSE, in line with previous reports, the latter displayed antiviral activity. Komura et al. demonstrated the efficacy of GSE as an antimicrobial agent on avian pathogens including avian influenza virus, Newcastle disease virus, infectious bursal disease virus, *Salmonella Infantis*, and *Escherichia coli* (Komura et al., 2019). Also, GSE has shown similar antiviral activities against human enteric pathogens including Hepatitis A virus in a dose-dependent manner (Su and D’Souza, 2011). Interestingly, GSE antiviral activity seems to be particularly effective on enveloped viruses. Since SARS-CoV-2 is an enveloped virus the GSE characteristics to induced or target the viral envelope should not be overlooked as candidate therapies for COVID-19 emerge (Schoeman and Fielding, 2019). On the other hand, xylitol did not show *in vitro* virucidal properties in the present study. However, it seems that the viral protective effects of xylitol are evident *in vivo* a suggested by studies demonstrating ameliorating effects against human respiratory syncytial virus and changes in the microbiota when consumed orally (Uebanso et al., 2017; Xu et al., 2016).

The precise mechanism of action of GSE is poorly understood. However, according to the present virucidal tests, the active component of the spray is the GSE, which is in line with previous reports demonstrating that the extract was is effective to inactivate different enveloped and non-enveloped viruses (Su and D’Souza, 2011).

Moreover, it seems that the mechanism of action of GSE targets the viral adsorption (or viral binding) to a greater extent than viral replication. It is worth mentioning that studies of the precise mechanism of action of GSE are beyond the scope of this work.

As with any research study, the present experimental design is not free from some limitations. The minimum time required for the Xlear solution to exert the virucidal effect was not investigated. Furthermore, to assess the relevance of the time-dependent effect of Xylitol effect *in vivo*, it will be important to verify if the addition of the spray-on cells previously infected at nontoxic doses would exert a reduction of the viral titer. Also, whether pre-treating the cells with the spray and subsequently adding the virus would decrease the rate of infection would be needed to assess the possible preventive use of the nasal spray.

## CONCLUSIONS

This study demonstrates the strong virucidal effects against SARS-CoV-2 of the Xlear nasal spray compound with xylitol and GSE. Using a virucidal nasal spray could become a cutting-edge element in the prevention and treatment of COVID-19 disease. To further ascertain the impact of this nasal spray in SARS-CoV-2, we propose to perform further a randomized placebo-controlled study of intranasally delivered Xlear in patients with mild to moderate SARS-CoV-2 and randomized placebo-controlled preventive trial in healthcare workers.

## Acknowledgments

The authors are grateful to Mr. Nathan Jones for donating the reagents and testing solutions for this study.

## Funding

The study was funded thanks to the financial support of the “Fondation privée des HUG” and the Carigest Foundation to CT.

## Notes

Declarations of interest: none

### Competing Interest Statement

The authors have declared no competing interest.

## REFERENCES

Akgül, Ö., Ak, A., Zorlu, S., Özdaş, D., Uslu, M., Çayirgan, D., 2020. Effects of short-term xylitol chewing gum on pro-inflammatory cytokines and Streptococcus mutans: a randomized, placebo-controlled trial. Int J Clin Pract, e13623.

Chacón, M.R., Ceperuelo-Mallafré, V., Maymó-Masip, E., Mateo-Sanz, J.M., Arola, L., Guitiérrez, C., Fernandez-Real, J.M., Ardèvol, A., Simón, I., Vendrell, J., 2009. Grape-seed procyanidins modulate inflammation on human differentiated adipocytes in vitro. Cytokine 47(2), 137–142.

Dong, E., Du, H., Gardner, L., 2020. An interactive web-based dashboard to track COVID-19 in real time. Lancet Infect Dis 20(5), 533–534.

Haukioja, A., Söderling, E., Tenovuo, J., 2008. Acid production from sugars and sugar alcohols by probiotic lactobacilli and bifidobacteria in vitro. Caries Res 42(6), 449–453.

Higgins, T.S., Wu, A.W., Illing, E.A., Sokoloski, K.J., Weaver, B.A., Anthony, B.P., Hughes, N., Ting, J.Y., 2020. Intranasal Antiviral Drug Delivery and Coronavirus Disease 2019 (COVID-19): A State of the Art Review. Otolaryngol Head Neck Surg, 194599820933170.

Hoffmann, M., Kleine-Weber, H., Krüger, N., Müller, M., Drosten, C., Pöhlmann, S., 2020. The novel coronavirus 2019 (2019-nCoV) uses the SARS-coronavirus receptor ACE2 and the cellular protease TMPRSS2 for entry into target cells. bioRxiv.

Jia, H.P., Look, D.C., Shi, L., Hickey, M., Pewe, L., Netland, J., Farzan, M., Wohlford-Lenane, C., Perlman, S., McCray, P.B., Jr., 2005. ACE2 receptor expression and severe acute respiratory syndrome coronavirus infection depend on differentiation of human airway epithelia. J Virol 79(23), 14614–14621.

Komura, M., Suzuki, M., Sangsriratanakul, N., Ito, M., Takahashi, S., Alam, M.S., Ono, M., Daio, C., Shoham, D., Takehara, K., 2019. Inhibitory effect of grapefruit seed extract (GSE) on avian pathogens. J Vet Med Sci 81(3), 466–472.

Meister, T.L., Brüggemann, Y., Todt, D., Conzelmann, C., Müller, J.A., Groß, R., Münch, J., Krawczyk, A., Steinmann, J., Steinmann, J., Pfaender, S., Steinmann, E., 2020. Virucidal Efficacy of Different Oral Rinses Against Severe Acute Respiratory Syndrome Coronavirus 2. J Infect Dis 222(8), 1289–1292.

Salli, K., Lehtinen, M.J., Tiihonen, K., Ouwehand, A.C., 2019. Xylitol’s Health Benefits beyond Dental Health: A Comprehensive Review. Nutrients 11(8).

Schoeman, D., Fielding, B.C., 2019. Coronavirus envelope protein: current knowledge. Virol J 16(1), 69.

Siddiqi, H.K., Mehra, M.R., 2020. COVID-19 illness in native and immunosuppressed states: A clinical-therapeutic staging proposal. J Heart Lung Transplant 39(5), 405–407.

Storey, D., Lee, A., Bornet, F., Brouns, F., 2007. Gastrointestinal tolerance of erythritol and xylitol ingested in a liquid. Eur J Clin Nutr 61(3), 349–354.

Su, X., D’Souza, D.H., 2011. Grape seed extract for control of human enteric viruses. Appl Environ Microbiol 77(12), 3982–3987.

Uebanso, T., Kano, S., Yoshimoto, A., Naito, C., Shimohata, T., Mawatari, K., Takahashi, A., 2017. Effects of Consuming Xylitol on Gut Microbiota and Lipid Metabolism in Mice. Nutrients 9(7).

Ur-Rehman, S., Mushtaq, Z., Zahoor, T., Jamil, A., Murtaza, M.A., 2015. Xylitol: a review on bioproduction, application, health benefits, and related safety issues. Critical reviews in food science and nutrition 55(11), 1514–1528.

Weissman, J.D., Fernandez, F., Hwang, P.H., 2011. Xylitol nasal irrigation in the management of chronic rhinosinusitis: a pilot study. Laryngoscope 121(11), 2468–2472.

Xu, M.L., Wi, G.R., Kim, H.J., Kim, H.J., 2016. Ameliorating Effect of Dietary Xylitol on Human Respiratory Syncytial Virus (hRSV) Infection. Biol Pharm Bull 39(4), 540–546.

